# Inhibition of UBE2N in regulatory T-cells boosts immunity against cancer

**DOI:** 10.1101/2024.10.22.619436

**Authors:** Wanying Miao, Vaibhav Jain, Madison Han, Yingai J. Jin, Georgia M. Beasley, Daniel T. Starczynowski, Simon G. Gregory, Jennifer Y. Zhang

## Abstract

Regulatory T (Treg) cells prevent autoimmunity and facilitate cancer immune evasion. Depletion of Tregs is a promising cancer therapy, but risks of autoimmune reactions hamper its clinical translation. Here, we demonstrate that temporally induced deletion of Ube2n in Tregs (Ube2n^Treg-KO^) of adult mice results in a robust expansion and activation of cytotoxic CD8^+^ T-cells in response to cancer cell challenges, producing a long-lasting survival benefit without autoimmune complications. The anti-tumor effect persists following adoptive T-cell transfer to T-cell-deficient Rag1-knockout mice. Single-cell transcriptomic analysis revealed that UBE2N deletion shifted immunosuppressive Tregs to effector-like T-cells. This shift is characterized by the downregulation of c-Myc target genes, resembling that observed in tumor-infiltrating Tregs of melanoma patients. Further analyses confirm that UBE2N maintains c-Myc protein stability via suppression of K48-Ubiquitin-mediated proteasomal degradation. Taken together, our studies uncover a hitherto unexplored and potentially druggable UBE2N/c-Myc signaling axis to eradicate Treg-enabled cancer immune escape.

## Introduction

Cancer cells employ a wide variety of strategies to grow and evade elimination by the immune system. Immune checkpoint blockade (ICB) with inhibitors of PD1/CTLA4 and respective ligands comprises the frontline immunotherapies for many advanced cancers, including melanoma ^1–4^. However, only 35-50% of patients with advanced melanoma show a prolonged life expectancy ranging from 1/2 to 4 years, with fewer surviving beyond the 5-year mark ^5,6^. Thus, there is an unmet need for effective treatments. In this regard, regulatory T-cells (Tregs) foster an immunosuppressive tumor microenvironment (TME) ^7–9^. Increased Tregs correlate with poor prognosis ^10,11^ and represent a major barrier to the success of immunotherapy ^12–14^. ICB therapies elevate cytotoxic effector T-cell function, but they can inadvertently increase Treg activity, leading to therapeutic resistance ^15–17^. Treg depletion is considered a promising cancer therapy ^18,19^. However, its clinical translation is hindered by the lack of Treg-specific depletion strategies and the risk of autoimmunity ^20,21^.

Tregs are characterized by the expression of IL-2 receptor α-chain (CD25) and Foxp3, a transcription factor constitutively expressed in Tregs to regulate gene expression, consequently maintaining the suppressive function and proliferation of Tregs ^22,23^. While Foxp3 is essential for Treg lineage specification, it is insufficient to confer Treg function ^24,25^. Other gene regulators are required at distinct stages of Treg differentiation, maintenance, and function ^26^. For example, KLF2 controls the migration of naïve Tregs to secondary lymphoid organs ^27^. c-Myc is required for Treg differentiation from naïve T-cells and conventional T-cells, but, interestingly, it is dispensable for the maintenance of existing mature Tregs ^28^. Understanding the complex regulatory network governing Treg function is crucial for rational targeted therapies.

UBE2N (also known as UBC13) is an E2 ubiquitin-conjugating enzyme that specifically catalyzes lysine 63-Ubiquitination (K63-Ub) to regulate signal transduction of the NF-κB and MAPK signaling pathways, gene expression, and DNA repair ^29,30^. Our previous study demonstrated that UBE2N promotes melanoma growth via MEK/FRA1/SOX10 signaling ^31^. Additionally, Foxp3^Cre^-mediated Treg-specific deletion of Ube2n in developing embryos leads to the early onset of autoimmunity during adulthood, which is attributed to decreased Treg function and conversion of Treg into Th1/Th17 CD4^+^ effector-like T-cells ^32^. Interestingly, neither the number of Tregs nor Foxp3 expression was reduced in these mutant animals, suggesting that UBE2N acts through regulation of other gene regulators to ensure proper Treg development and function. These findings underscore UBE2N as a potential cancer therapeutic target. However, the roles of UBE2N in Treg in the context of solid tumor growth have not been explored.

## Results

### Treg-targeted deletion of UBE2N induces a strong immunity against cancer

To test whether the deletion of Ube2n specifically in Tregs affects tumor growth, we crossed Foxp3^Cre-ERT2^ mice with Ube2n^fl/fl^ mice and utilized Foxp3^Cre-ERT2^.Ube2n^fl/fl^ (Ube2n^Treg-KO^) and Foxp3^wt^.Ube2n^fl/fl^ (WT) litters for syngeneic tumor growth analysis (Fig. 1a). We first treated animals with daily intraperitoneal injections of tamoxifen for 3 consecutive days and verified UBE2N deletion by RT-PCR with CD4^+^CD25^+^ T-cells isolated from the spleen 2 weeks after tamoxifen treatment. Consistent with previous reports ^33^, the efficiency of Foxp3^+^ Treg-specific UBE2N deletion was approximately 30% (Fig. 1b). As expected, tumors in WT animals exhibited linear growth, reaching the humane endpoint by day 18. In contrast, tumors in Ube2n^Treg-KO^ grew significantly slower, and a majority of them became invisible by day 18 (Fig. 1c-d). About 60% of Ube2n^Treg-KO^ mice survived and appeared healthy beyond day 30 (Fig. 1e). Body weight loss occurred in WT mice due to exponential tumor growth, and it was not evident in Ube2n^Treg-KO^ mice (Fig. S2a). Additionally, the observed antitumor effect is applicable to the syngeneic MC38 colon cancer model (Fig. 1f).

**Fig. 1:**
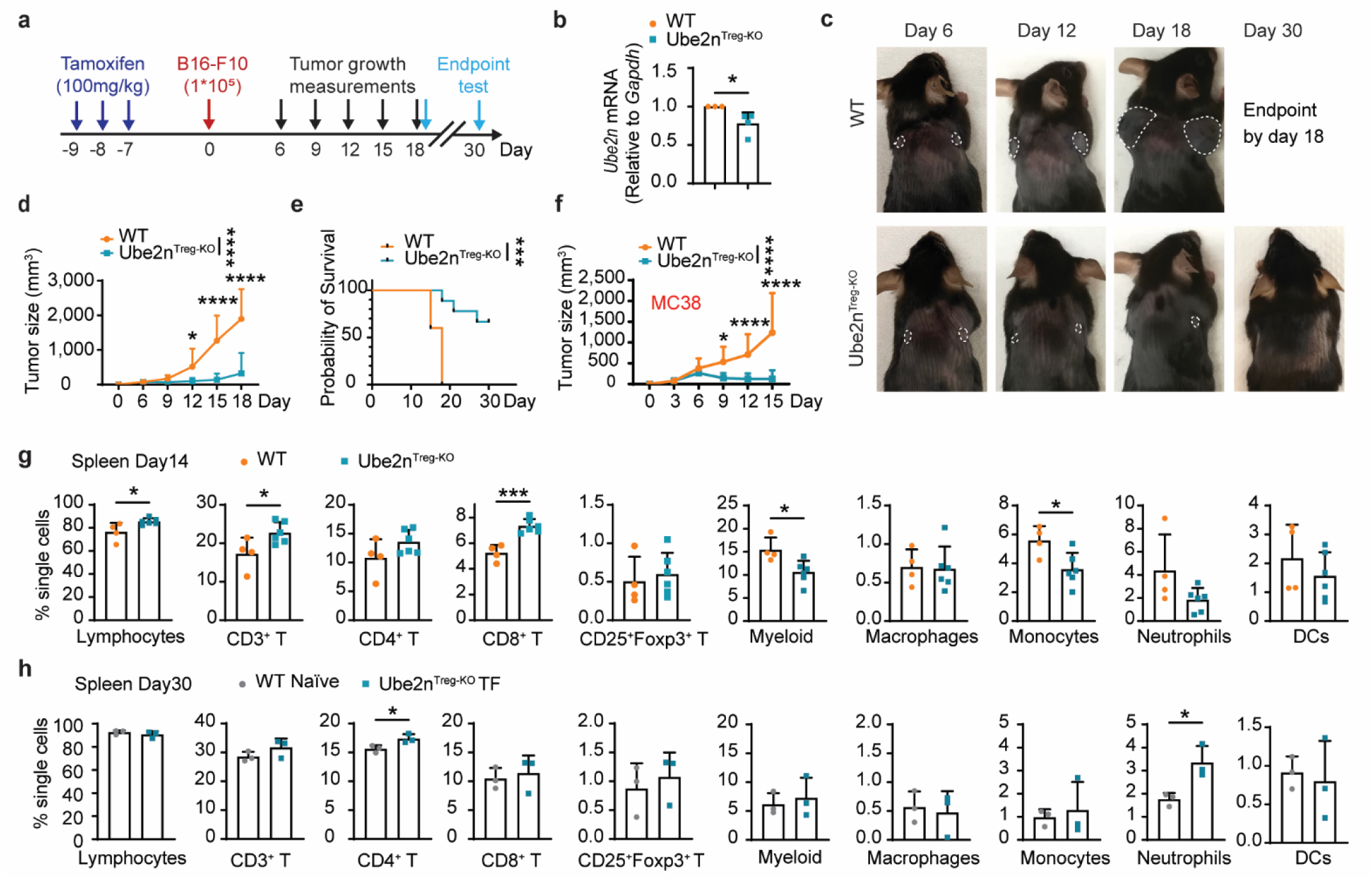
Treg-specific deletion of UBE2N inhibits tumor growth. **a**, Diagram of the experiment timeline. **b**, mRNA expression level of Ube2n in Tregs collected from the spleen 2 weeks after tamoxifen injection. *n* = 3-4 mice / group, *t* - Test. **c**, Representative image of animals with B16-F10 tumors. **d**, Quantification of tumor size. WT *n* = 10, Ube2n^Treg-KO^ *n* = 18 tumors, Two-way ANOVA and Bonferroni. **e**, Survival rate of tumor-bearing mice 30 days after tumor implantation. WT *n* = 5, Ube2n^Treg-KO^ *n* = 9 mice, Log-rank test. **f**, Quantification of MC38 tumors. WT *n* = 12, Ube2n^Treg-KO^ *n* = 16 tumors, Two-way ANOVA and Bonferroni. **g-h**, Quantification of spleen immune cells 14 days (**g**) and 30 days (**h**) after tumor implantation. Day 14, *n* = 4-6 mice / group, *t* - Test. Day 30, *n* = 3 mice / group, *t* - Test. Data are shown as mean ± SD unless otherwise specified. *p < 0.05, **p < 0.01, ***p < 0.001, ****p < 0.0001.

### UBE2N-deletion in adult Tregs boosts peripheral immune responses to tumor cells with no signs of autoimmune reactions

Foxp3^Cre^-mediated deletion of Ube2n during embryonic development results in the early onset of autoimmunity in adult mice ^32^. We asked whether Treg-specific deletion of UBE2N during adulthood affects the systemic immune environment. To address this question, we first isolated splenocytes from non-tumor bearing naïve animals 2 weeks after tamoxifen induction and performed flow cytometry (Fig. S1a and S1b). At a gross level, spleen and body weights of Ube2n^Treg-KO^ mice were comparable to the counterparts of WT mice (Fig. S1c). Flow cytometry revealed no significant differences in spleen immune cell populations between WT and Ube2n^Treg-KO^ mice (Fig. S1d), indicating that Treg-specific deletion of Ube2n does not affect the peripheral immune cell populations in naïve adult mice. Next, we asked if UBE2N-deletion in Tregs affects peripheral immune responses in the context of tumor growth. We evaluated immune cells of the spleen and blood 14 days after tumor implantation . The spleen weights of WT and Ube2n^Treg-KO^ tumor-bearing mice were indifferent (Fig. S2b). However, the percentage of lymphocytes significantly increased, while myeloid cells significantly decreased in the spleen of Ube2n^Treg-KO^ mice compared to WT counterparts. Further evaluation of the lymphocyte subpopulations revealed an increase of CD3^+^ T-cells and CD8^+^ T-cells in Ube2n^Treg-KO^ mice. Monocytes were decreased in Ube2n^Treg-KO^ mice, whereas macrophages, neutrophils, and dendritic cells (DCs) were not significantly affected by UBE2N-loss in Tregs (Fig. 1g). Similarly, blood samples from Ube2n^Treg-KO^ tumor-bearing mice had increased T-lymphocyte and decreased myeloid cell populations (Fig. S2d). Interestingly, the percentages of Tregs in both spleen and blood samples were comparable between WT and mutant animals (Fig. 1g and Fig. S2d). These results indicate that depletion of UBE2N in Tregs increases peripheral immune responses in the context of tumor growth without impairing Treg survival.

Next, we asked whether the peripheral immune response persists after tumor eradication. We examined the spleen, blood, and thymus of tumor-free Ube2n^Treg-KO^ mice (Ube2n^Treg-KO^ TF) 30 days after tumor implantation. The body and spleen weights of Ube2n^Treg-KO^ TF mice were comparable to their age-matched naïve WT counterparts (WT B16^-^) (Fig. S2c). The percentages of spleen lymphocytes, CD3^+^ T-cells, CD8^+^ T-cells, Foxp3^+^ T-cells, myeloid cells, macrophages, monocytes, and DCs did not show significant differences between groups. Percentages of neutrophils were increased from 1.8% to 3.3%, and there was a slight increase of CD4^+^ T-cells in the Ube2n^Treg-KO^ TF group (Fig. 1h). Immune cell profiles of the blood and thymus were also comparable between the two groups (Fig. S2e-g), with only DCs showing a slight decrease in the blood of Ube2n^Treg-KO^ TF group (Fig. S2e). These results indicate that UBE2N-deletion in Tregs boosts antitumor immunity without incurring a prolonged autoimmune response after tumor eradication.

### Treg-specific deletion of UBE2N exerts a persistent anti-tumor effect

To investigate whether UBE2N deficiency in Tregs induces a prolonged anti-tumor effect, we subjected Ube2n^Treg-KO^ TF mice to a second injection of B16-F10 cells 78 days after the first challenge (Fig. 2a). Again, our data show that 55.6% of Ube2n^Treg-KO^ mice were tumor-free (TF) 30 days after the first injection (Fig. 2b). A majority of Ube2n^Treg-KO^ TF animals showed a persistent resistance to tumor growth following the second round of cancer cell challenge (Fig. 2c). These data indicate that temporally induced Ube2n deletion in Tregs renders a majority of animals a long-term immunity against tumor growth.

**Fig. 2:**
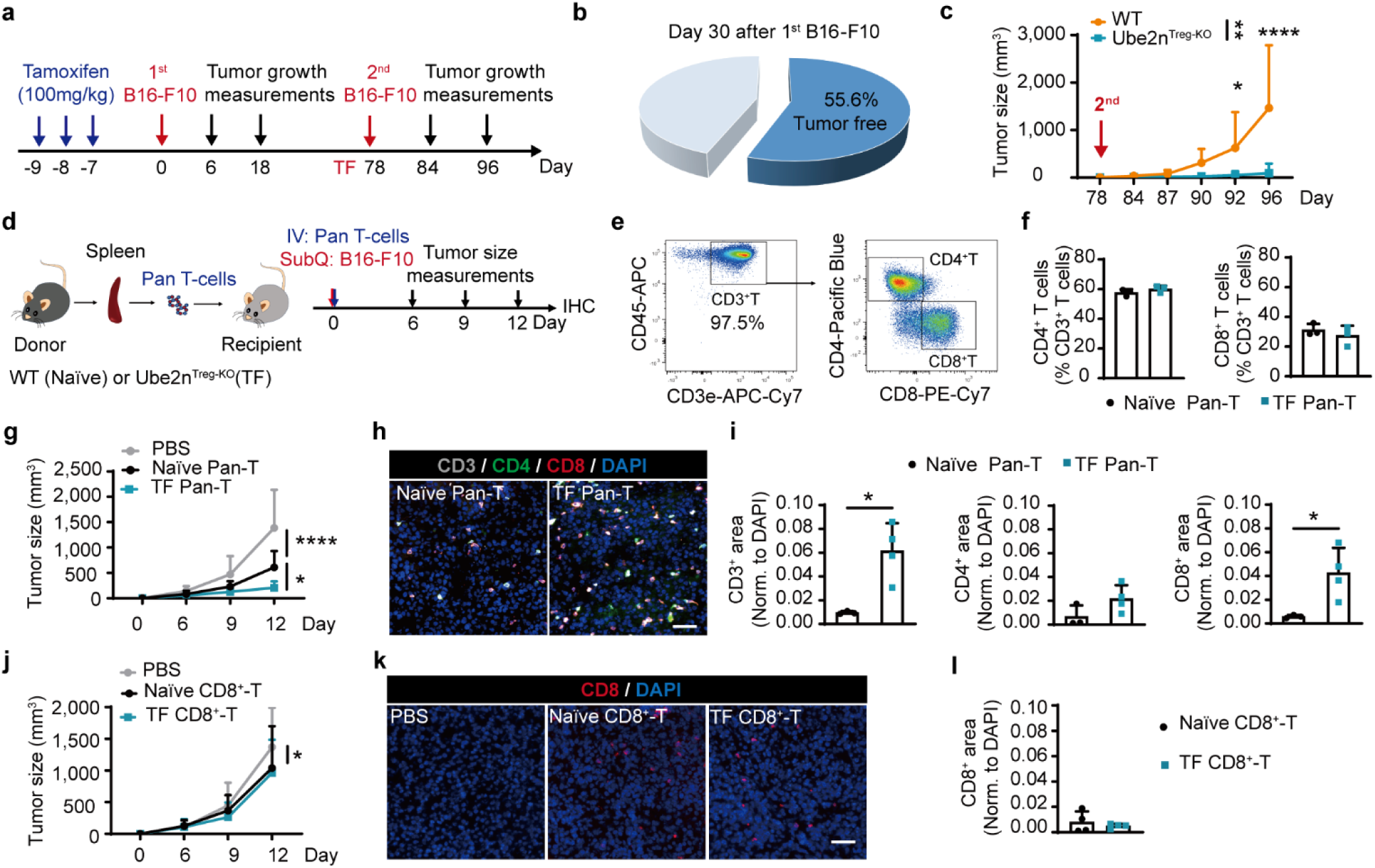
Long-lasting anti-tumor effects of temporally induced UBE2N deletion in Tregs. **a**, Diagram of the experiment timeline. **b**, Quantification of tumor-free (TF) Ube2n^Treg-KO^ mice on day 30 after the 1^st^ injection. Ube2n^Treg-KO^ mice with palpable *n* = 12, TF *n* = 15. **c**, Quantification of tumor size after the 2^nd^ challenge. WT *n* = 6, Ube2n^Treg-KO^ *n* = 12 tumors, Two-way ANOVA and Bonferroni. **d**, Diagram of the adoptive T-cell transfer (ACT). Spleen pan-T-cells isolated from naïve wild-type (WT) or TF Ube2n^Treg-KO^ mice were intravenously injected (1x10^6^ cells) into Rag1-KO mice within 10 min after subcutaneous injection of B16-F10 cells (1x10^5^ cells). Tumor size was measured 6, 9, and 12 days post-tumor implantation. **e**, Gating strategy of donor spleen Pan-T-cells. **f**, Quantification of percentages of CD4^+^ and CD8^+^ T-cells among CD3^+^ T-cells. *n* = 3 / group, *t* - test. **g**, Quantification of B16-F10 tumor size after ACT of Pan-T-cells. PBS *n* = 14, Naïve Pan-T *n* = 14, TF Pan-T *n* = 16 tumors, Two-way ANOVA and Bonferroni. **h**, Representative immunostaining of CD3 (gray), CD4 (green), and CD8 (red), DAPI nuclear staining (blue) in tumors collected 12 days after ACT of Pan-T-cells. Scale bar = 50 μm. **i**, Quantification of CD3, CD4, and CD8 signal positive area. Naïve Pan-T *n* = 3 tumors, TF Pan-T *n* = 4 tumors, *t* - test. **j**, Quantification of the B16-F10 tumor size after CD8^+^ T cell treatment. PBS *n* = 14, Naïve Pan-T *n* = 12, TF Pan-T *n* = 14 tumors, Two-way ANOVA and Bonferroni. **k**, Representative immunostaining of CD8^+^ T-cells (red), and DAPI nuclear staining (blue) in tumors collected 12 days after CD8^+^ T-cell treatment. Scale bar = 50 μm. **l**, Quantification of CD8 signal positive area. *n* = 4 tumors / group, *t* - Test. Data are shown as mean ± SD unless otherwise specified. *p < 0.05, **p < 0.01, ***p < 0.001, ****p < 0.0001.

Next, we tested whether the anti-tumor effect of UBE2N-deficient Tregs is mediated by T-lymphocytes and transferrable. We performed intravenous adoptive cell transfer (ACT) of pan-T-cells from Ube2n^Treg-KO^ TF mice to Rag1^-/-^ recipient mice which otherwise lack endogenous mature T-cells and hence support allografts (Fig. 2d). T-cells were isolated from the spleens of age-matched naïve WT mice (naïve pan-T) and Ube2n^Treg-KO^ TF mice (TF pan-T). The purity (∼97.5%) and the percentages of CD4^+^ and CD8^+^ T-cells were comparable between the two groups (Fig. 2e-f). Compared to PBS control, both naïve and TF Pan-T-cell treatments inhibited tumor growth, though the latter exhibited a significantly enhanced antitumor effect (Fig. 2g). Immunostaining of the tumor tissues revealed an over 6-fold increase of tumor-infiltrating CD8^+^ T-cells and a small but insignificant increase of CD4^+^ T-cells in the TF pan-T group (Fig. 2h-i). To further determine whether CD8^+^ T-cells are sufficient to retain the antitumor memory, we performed ACT of CD8^+^ T-cells isolated from the spleens of naïve WT and Ube2n^Treg-KO^ TF mice. Interestingly, compared to PBS control, both naïve WT and TF CD8^+^ T-cells decreased tumor growth by about 25% at day 12, but the two groups showed no difference in both tumor growth and infiltration of CD8^+^ T-cells (Fig. 2j-l). Together, these results indicate that the Ube2n^Treg-KO^-driven antitumor effect is retained after ACT and requires the presence of CD4^+^ T-cells to sustain CD8^+^ T-cell expansion.

### Treg-specific deletion of UBE2N increases tumor-infiltrating CD8^+^ T-cells in existing tumors

To further determine how UBE2N deletion in Tregs affects the existing tumor microenvironment, we subjected animals first to subcutaneous injection of B16-F10 cells and then to intraperitoneal injections of tamoxifen every 2-3 days until the endpoint (Fig. 3a). Data shows that tumors grew significantly slower in Ube2n^Treg-KO^ mice than in WT mice (Fig. 3b), which was correlated by significant increases of CD45^+^ and CD8^+^ immune cells in the mutant tumors. (Fig. 3c-e). In contrast, percentages of CD4^+^ T-cells were similar between WT and Ube2n^Treg-KO^ tumors (Fig. 3e). Further flow cytometry verified that while CD45^-^ non-immune cells were decreased in Ube2n^Treg-KO^ tumors compared to WT tumors, CD45^+^ immune cells, CD3^+^, and CD8^+^ T-cells were increased, as were CD11b^+^ myeloid lineage cells, including monocytic-myeloid derived suppressor cells (M-MDSC), and DCs (Fig. 3f). Interestingly, CD4^+^ T-cells, Foxp3^+^ Tregs, and neutrophil populations were comparable between WT and mutant tumors. Consistent with a highly immune active TME, percentages of IFN-γ^+^CD8^+^ T-cells and overall IFN-γ expression were significantly higher in mutant tumors than their WT counterparts (Fig. 3g). Additionally, analysis of the mutant tumors of varying sizes showed that the relative numbers of IFN-γ^+^CD8^+^ T-cells were inversely correlated with tumor sizes (Fig. 3h). RT-PCR results showed that proinflammatory cytokines including Tnfa, Ifng, and Il6, as well as Il10 and Tgfb increased in mutant tumors (Fig. 3i). Together, these results indicate that Treg-targeted deletion of Ube2n boosts immune activities in the established TME.

**Fig. 3:**
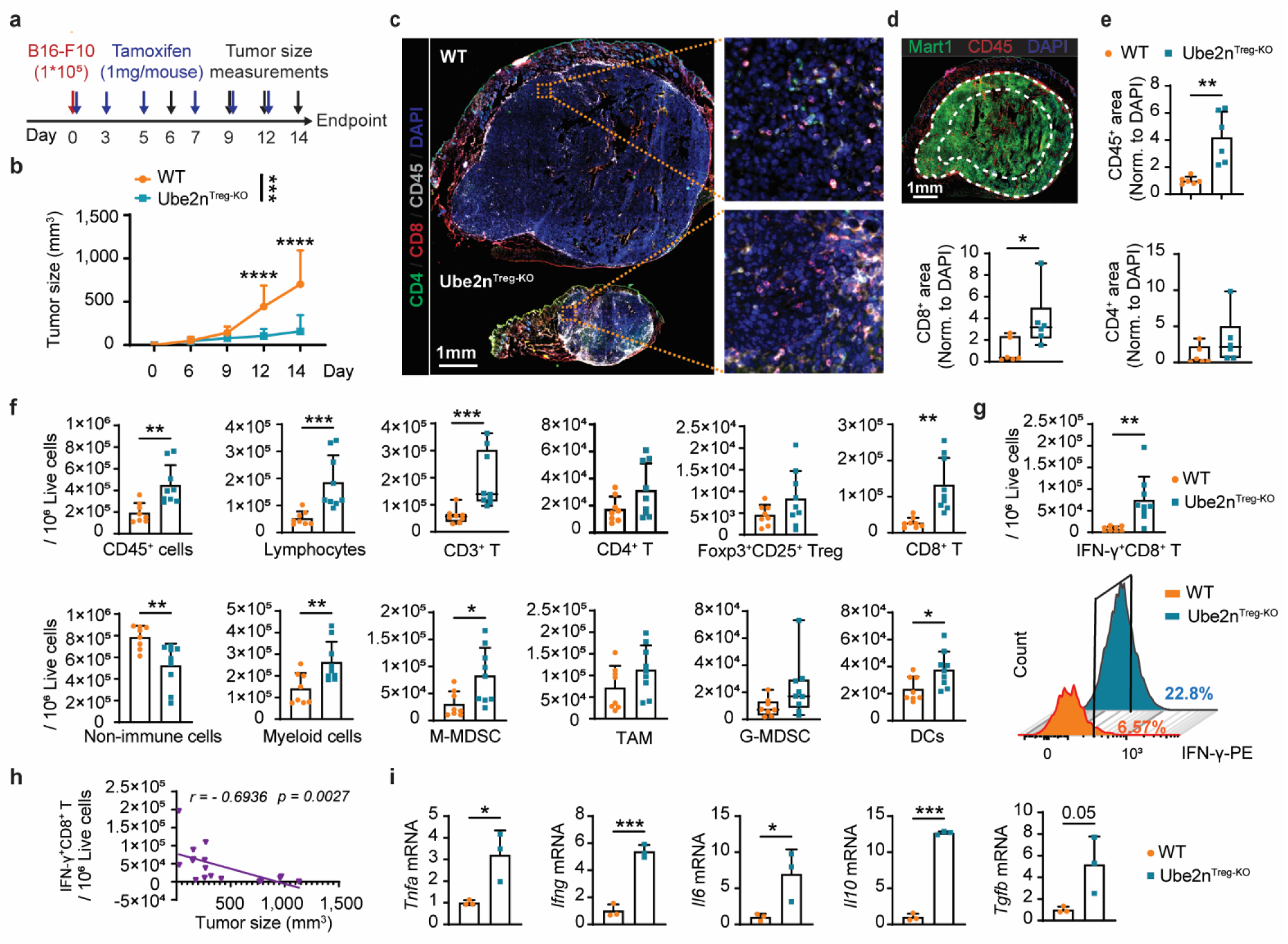
Treg-specific deletion of Ube2n in mice with preexisting melanoma cells inhibits tumor growth and increases immune cell infiltration. **a**, Diagram of the experiment timeline. **b**, B16-F10 tumor size. WT *n* = 14 tumors, Ube2n^Treg-KO^ *n* = 16 tumors, Two-way ANOVA and Bonferroni. **c**, Representative immunostaining image of T cell markers CD4 (green), CD8 (red), immune cell marker CD45 (gray), and DAPI nuclear staining (blue) in the tumors 14 days after implantation. **d**, Representative immunofluorescence images indicating the area for quantification (between white dotted lines). Mart1 (green), CD45 (red), and DAPI (blue). Scale bar = 1 mm. **e**, Quantification of infiltrating immune cells within 1 mm of the tumor boundary. CD45, CD8, and CD4 signal positive areas were normalized to DAPI, *n* = 6 tumors / group, *t* - Test. **f**, Quantification of immune cells (CD45^+^), non-immune cells (CD45^-^), lymphocytes (CD45^+^CD11b^-^), CD3^+^ T-cells, CD4^+^ T-cells, CD8^+^ T-cells, Tregs, myeloid cells (CD45^+^CD11b^+^), granulocytic-myeloid derived suppressor cells (G-MDSC: CD11b^+^CD11c^-^Ly6G^+^F4/80^-^), monocytic-myeloid derived suppressor cells (M-MDSC: CD11b^+^CD11c^-^Ly6G^-^F4/80^-^Ly6C^+^), tumor-associated macrophage (TAM: CD11b^+^CD11c^-^Ly6G^-^ F4/80^+^) and DCs 14 days after tumor implantation. *n* = 8-9 tumors / group, *t* - Test. **g**, Quantification of IFN-γ^+^CD8^+^ T-cells in day 14 tumors (upper). Histograms show examples of the percentage of IFN-γ^+^CD8^+^ T-cells among CD8^+^ T-cells (bottom). **h**, Spearman correlation analysis between tumor size and the number of IFN-γ^+^CD8^+^ T-cells. *n* = 8-9 tumors/group, *t* - Test. **i**, mRNA expression levels of cytokines in day 14 tumors. *n* = 3 tumors / group, *t* - Test. Data are shown as mean ± SD unless otherwise specified. (**e**: CD8^+^ and CD4^+^; **f**: CD4^+^ T-cells and G-MDSC; Data are presented as a box-and-whisker plot, with bounds from 25^th^ to 75^th^ percentile). *p < 0.05, **p < 0.01, ***p < 0.001.

### scRNA analysis revealed an enhanced cytotoxic CD8^+^ T cell expansion in the TME

To elucidate the molecular mechanisms underlying the antitumor immune response of UBE2N-deficient Tregs, we isolated CD4^+^ and CD8^+^ tumor-infiltrating lymphocytes (TILs) 14 days post-tumor implantation and tamoxifen treatment. A total of 9,120 WT and 8,621 Ube2n^Treg-KO^ TILs were collected from 3 animals per group and subjected to 10Xgenomic single-cell transcriptome analyses. UMAP analysis identified 22 clusters, with over 90% of these cells being CD4^+^ and CD8^+^ T-cells. Based on the differentially expressed gene (DEG) markers, cells were identified as CD8^hi^ T-cells, CD8^lo^ T-cells, CD4^+^ T-cells, DCs (Itgax, Btla, Lilrb4a, Nrp1, Zbtb46), natural killer cells (Klrb1a), macrophages (Itgam, Adgre1), and melanoma cells (Fig. 4a and Fig. S3a). The percentage of CD8^hi^ T-cells increased in the Ube2n^Treg-KO^ sample compared to WT (55.9% Vs. 34.0%)), while the percentage of CD8^lo^ T-cells decreased (10.5% Vs. 39.8%), CD4^+^ T-cells increased in Ube2n^Treg-KO^ (23.0% Vs. 11.8%), but CD4^+^ T-cells represented a small fraction compared to CD8^+^ T-cell populations (Fig. 4b). In agreement with the increased abundance of CD8^+^ T-cells, Ifng, a marker that positively correlates with the anti-tumor effect of cytotoxic T-cells, showed increased expression in the Ube2n^Treg-KO^ group, predominantly in CD8^+^ T-cell clusters (Fig. 4c). To further investigate the functional changes of CD8^+^ T-cells, we subclustered CD8^+^ T-cells into effector, effector/exhaustion, and naïve/memory subtypes based on marker expressions ^34^. CD8^+^ effector T-cells (cluster 2) expressed high levels of effector marker genes, including Prf1, Fasl, Gzmb, Ifng, Ccl5, and Ccl3, and low levels of exhaustion marker genes, including Lag3, Pdcd1, Tigit, Havcr2, Entpd1, and Tox. CD8^+^ effector/exhaustion T-cells (clusters 0 and 14) had high expression of exhaustion markers. Naïve/memory CD8^+^ T-cells (cluster 9) expressed high levels of Sell, Il7r, Lef1, Ccr7, and Tcf7. Proliferating CD8^+^ T-cells (clusters 3, 17, and 4) had high expression of Mki67. Inactive CD8^+^ T-cells (clusters 11, 10, 12, 13, 18, and 20) had low-level expression of Cd8a and other marker genes (Fig. 4d). The percentages of effector (20.9% Vs. 6.8%) and effector/exhaustion CD8^+^ T-cells (34.3% Vs. 15.9%), as well as naïve/memory CD8^+^ T-cells (9.2% Vs. 4.1%) were increased in the Ube2n^Treg-KO^ group versus the WT group, while inactive CD8^+^ T-cells dramatically decreased (15.8% Vs. 53.9%) (Fig. 4e). To evaluate functional differences between WT and mutant groups, we performed gene ontology (GO) analysis of DEGs of cluster 0 (effector/exhaustion CD8^+^ T-cells) (Fig. S3b upper panel). The top relevant terms, including interferon-mediated signaling pathway, CD8^+^ alpha-beta T-cell activation, and positive regulation of NK cell-mediated immunity, were significantly enriched in cluster 0 of the Ube2n^Treg-KO^ group (Fig. 4f). Similar results were shown in cluster 2 (effector CD8^+^ T-cells) (Fig. S3b bottom panel and S3c). Taken together, these results confirm that deletion of UBE2N in Tregs enhances cytotoxicity CD8^+^ T-cell activity in the TME.

**Fig. 4:**
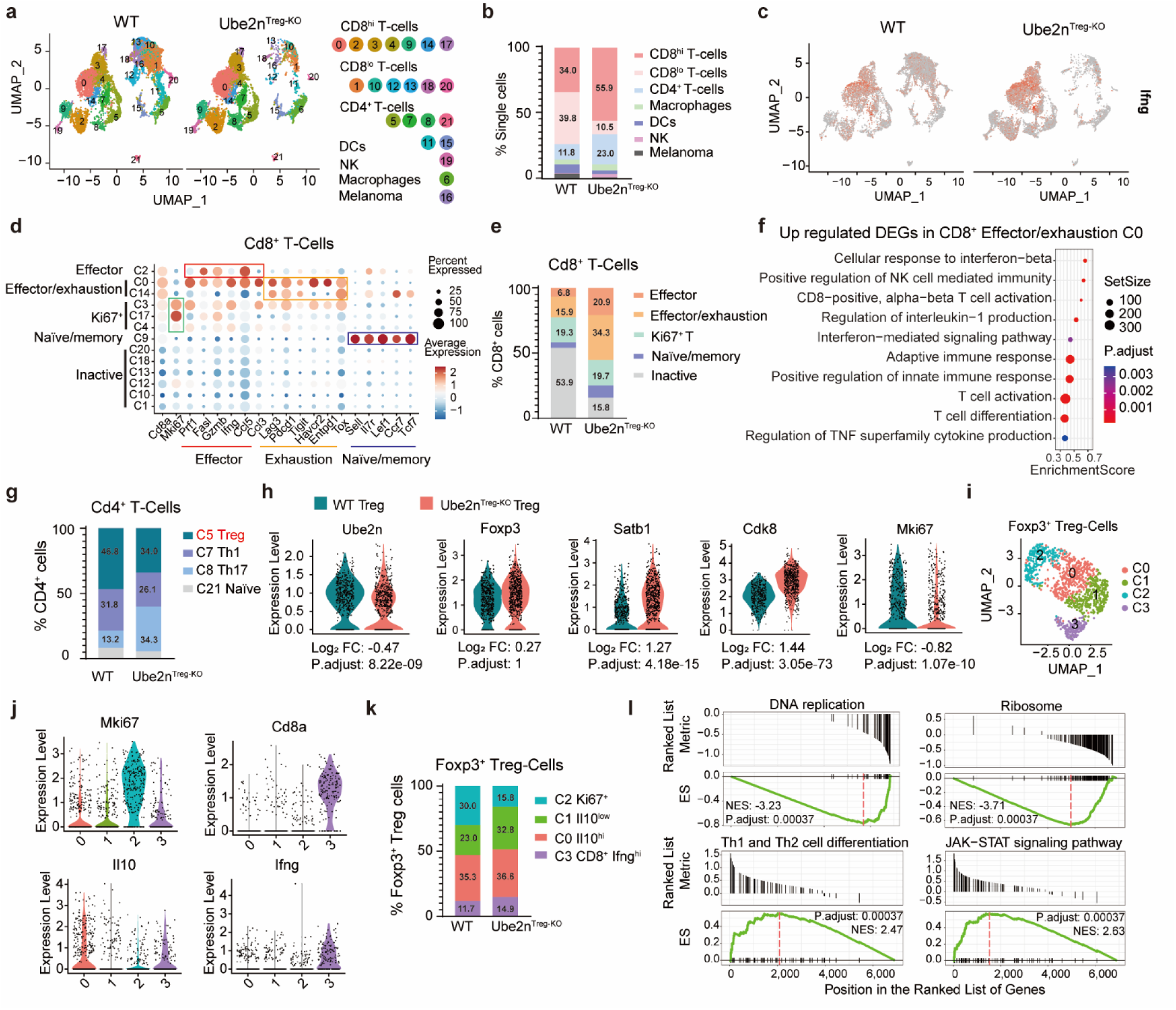
Single-cell transcriptome analysis reveals dysfunctional Tregs accompanied by increased cytotoxic T-cell expansion and activation in Ube2n^Treg-KO^ tumors. **a**, UMAP depicting 22 clusters of 9120 and 8621 tumor-infiltrating T-cells derived from WT and Ube2n^Treg-KO^ B16-F10 tumors, respectively. **b**, Percentages of immune cell populations for each group. **c**, UMAP plot Ifng gene expression profile in WT and Ube2n^Treg-KO^ group. **d**, Dot plots indicating average expression of a panel of marker genes (x-axis) associated with effector, exhaustion, naïve/memory, and Ki67^+^ T-cell phenotypes for the Cd8^+^ T-cell clusters (y-axis). **e**, Percentages of CD8^+^ T-cell subpopulations for each group. **f**, GO enrichment of upregulated DEGs of WT vs. Ube2n^Treg-KO^ in CD8^+^ effector/exhaustion cells (cluster 0). **g**, Percentages of CD4^+^ T-cell subpopulations. **h**, Violin plots of marker gene expression in Tregs separated by groups. **i**, UMAP of integrated 1177 Tregs from WT and Ube2n^Treg-KO^ samples. **j**, Violin plots of marker gene expression in Tregs subpopulations. **k**, Percentages of Treg subpopulations for each group. **l**, Gene-set enrichment analysis (GSEA) of DNA replication, ribosome, Th1 and Th2 cell differentiation, and JAK-STAT signaling pathway response signatures in Tregs from WT compared to Ube2n^Treg-KO^. NES = Normalized Enrichment Score.

### UBE2N-deficient Tregs show dysfunction and low proliferation in the TME

Next, we investigated how UBE2N loss affects Tregs and CD4^+^ T-helper (Th) cells in the TME. Tregs (cluster 5) were identified based on the expression of Foxp3, Il10, and IL2ra (CD25). Th1 cells (cluster 7) had high expression of Tbx21, Ifng, and Tnf; Th17 (cluster 8) expressed Il17a, Tgfbr2, Rorc, and Il6ra; and naïve T-cells (cluster 21) expressed Sell and Il7r in addition to low-level Gzmb and Gata3 (Fig. S4a). The percentage of Tregs over the total CD4^+^ T-cells population was decreased in the Ube2n^Treg-KO^ group versus (34% Vs. 46.8%); Th17 cells increased (34.3% Vs. 13.2%), while Th1 cells slightly decreased (26.1% Vs. 31.8%) in Ube2n^Treg-KO^ versus the WT counterparts (Fig. 4g). Violin plot gene expression analysis confirmed that Ube2n was significantly decreased in Ube2n^Treg-KO^ Tregs (Fig. 4h). Ube2n loss did not affect Foxp3 expression but instead increased Satb1 and Cdk8, and decreased the cell proliferation marker Mki67 in Tregs (Fig. 4h). Satb1 is a transcription factor and chromatin organizer known to suppress Treg differentiation ^35^. Likewise, Cdk8 inhibits Treg differentiation ^36^. These results indicate that Treg differentiation is impaired in Ube2n^Treg-KO^ tumors.

To further investigate the functional changes of UBE2N-deficient Tregs, Foxp3^+^ Tregs (cluster 5) were subjected to further high-resolution UMAP analysis (Fig. 4i). Four clusters (C0-C3) were characterized by high expression of Mki67 (C2), Il10 (C0), Cd8a, and Ifng (C3) (Fig. 4j). IL-10 is secreted by Tregs and is often used as a surrogate marker to reflect the suppressive function of Tregs. The percentage of Il10^low^ Tregs (C1) increased, whereas the Mki67^+^ proliferating Tregs (C2) decreased in Ube2n^Treg-KO^ tumors (Fig. 4k). KEGG (Kyoto Encyclopedia of Genes and Genomes) analysis of DEGs (Fig. S4b) showed that DNA replication and ribosome pathways were suppressed in Ube2n^Treg-KO^ Tregs, while Th1 and Th2 cell differentiation and JAK-STAT signaling pathways were activated (Fig. 4l and Fig. S4c). Taken together, these data indicate that UBE2N-deletion in Tregs resulted in decreased proliferation and abnormal Treg differentiation and function.

### c-Myc transcription program is decreased in TIL Tregs Ube2n^Treg-KO^ mice and melanoma patient responders to immunotherapy

To characterize downstream effectors of UBE2N, we performed ChIP-X enrichment analysis (ChEA) on downregulated DEGs of Tregs using Enrichr ^37–39^. The bar graph shows that 4 of the top 10 relevant terms are Myc ChIP datasets (Fig. 5a), suggesting that UBE2N-loss repressed c-Myc-mediated transcription. To further investigate whether c-Myc-mediated transcription is relevant to tumor-infiltrating Tregs (TITs) of melanoma patients, we analyzed publicly available single-cell RNAseq data (GSE120575) of CD45^+^ immune cells derived from melanoma biopsies collected before and post treatments of PD1 and CTLA4 inhibitors ^40^. High-resolution UMAP of the original cluster 7 containing a total of 1,740 TITs were separated into 5 sub-clusters (Fig. 5b). Sub-cluster 1, which expresses classical markers of Treg (CD4, FOXP3, and CD25), was selected for further DEG and Enrichr analysis comparing responder (R) to non-responders (NR) of pre- and post-treatment samples (Fig. 5c). ChEA of the downregulated genes revealed Myc-mediated transcription being among the most significantly suppressed programs in pre-R TITs (161 cells of 9 patients) Vs. pre-NR TITs (87 cells of 10 patients) (Fig. 5d). A heatmap reveals that the expression levels of putative Myc target genes, such as HMGB2, HSBP1, HSPE-MOB4, PDCL3, GK, GEM, and NR3C1, are significantly decreased in pre-R TITs (Fig. 5e). Cross-comparisons of DEGs of mouse (Ube2n^Treg-KO^ Vs. WT) and patient (pre-R Vs. pre-NR) TITs revealed 152 overlapping genes (119 downregulated and 33 upregulated) (Fig. 5f). Gene enrichment analysis of these overlapping genes indicates increased T-cell activation and immune system processes (Fig. 5g) and decreased cell cycle, cell division, and protein kinase activities (Fig. 5h). The ChEA result shows that Myc-mediated transcription, along with that of FOXM1 and E2F1, is among the top terms associated with the overlapping genes (Fig. 5i). However, Myc mRNA expression was not decreased in both Ube2n-Ko mouse TITs and pre-R human TITs (Fig. 5j). These results indicate that decreased c-Myc activity in TITs is associated with a decreased Treg function and a better outcome in murine and human melanomas, and suggest that c-Myc is regulated at the protein level by UBE2N.

**Fig. 5:**
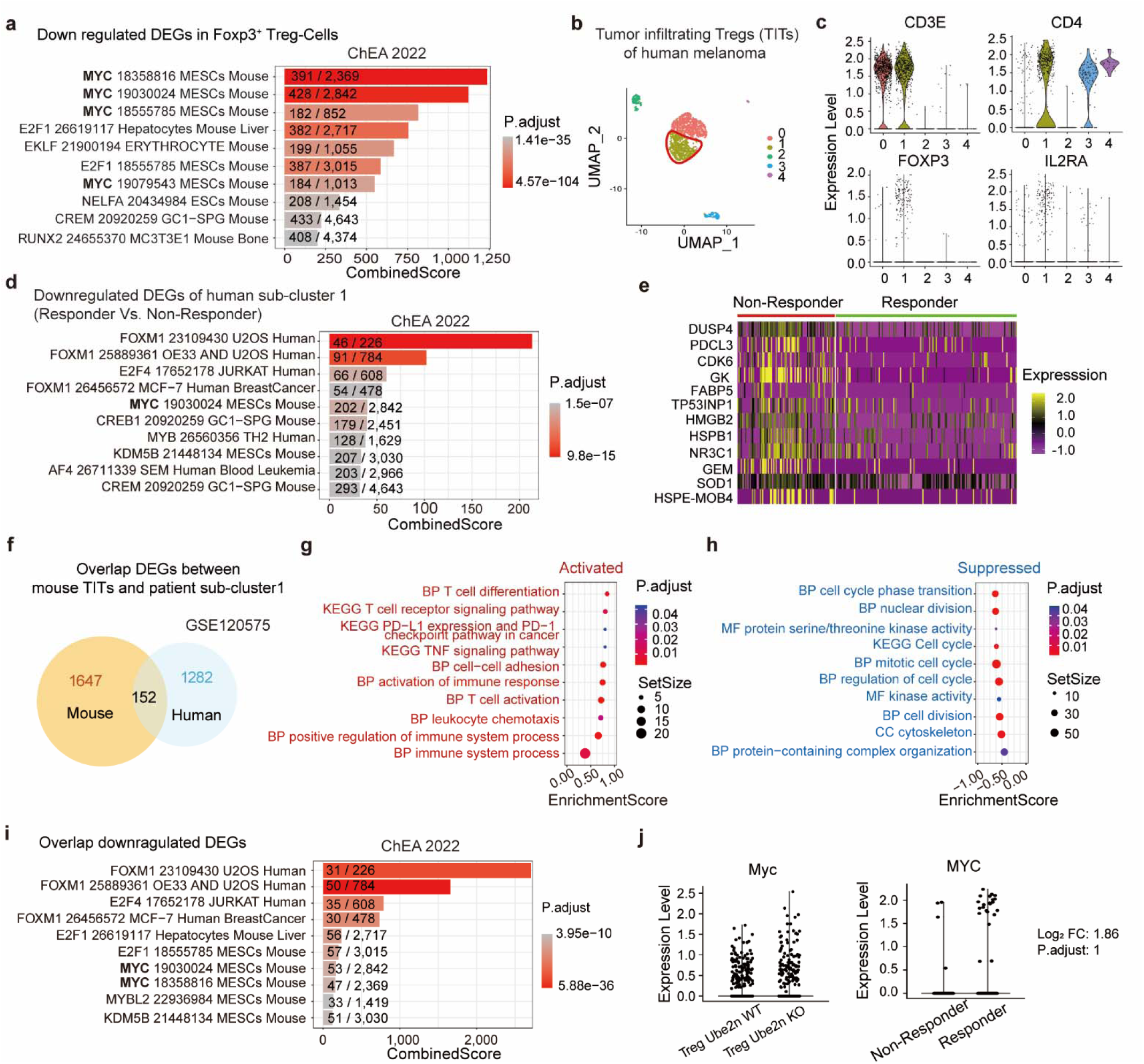
Decreased c-Myc activity in Ube2n-deficient Treg and tumor-infiltrating Tregs (TITs) of human patients. **a**, ChIP-X Enrichment Analysis (ChEA) of downregulated signatures in Tregs from WT compared with Ube2n^Treg-KO^. Numbers of matching genes are labeled on each bar. **b**, High resolution UMAP subclustering of 1740 TITs of the initial cluster 7 of single-cell RNAseq data acquired from 48 melanoma patients pre and post-treatments with PD1 and CTLA4 inhibitors (GSE120575). **c**, Violin plots of marker gene expression in TITs subpopulations. **d**, ChEA of downregulated DEGs in Sub-cluster 1 from pretreatment responder (Pre-R) Vs. non-responder (Pre-NR) patients. **e**, Heatmap of top c-Myc target genes that reached adjusted p.values < 0.05. **f**, The overlapping DEGs between mouse TITs and patients Sub-cluster 1. **g-h**, Gene enrichment results of activated (33 upregulated genes, **g**) and suppressed (119 downregulated genes, **h**) terms of overlapping genes. **i**, ChEA of overlapping downregulated DEGs searched against human and mouse ChiIP datasets. **j**, Myc expression level in mouse TITs (left) and patients Sub-cluster 1 (right).

### UBE2N regulates c-Myc stability in Treg

To test this possibility, we utilized the induced Treg (iTreg) model ^41,42^, which allowed us to obtain sufficient numbers of Tregs for in vitro analysis. Naïve CD4^+^ T-cells isolated from Rosa^CreERT2^.Ube2n^fl/fl^ splenocytes were treated with murine TGF-β/IL-2 and CD3/CD28 beads for 4 days to induce the iTreg phenotype, along with 4-hydroxytamoxifen (4-OHT) to induce Ube2n deletion (Fig. 6a). As expected, the mRNA levels of Foxp3 were significantly increased after 4 days of TGF-β/IL-2 treatment (Fig. 6b). IL-10 was decreased at both mRNA and protein levels in response to 4-OHT-induced deletion of UBE2N, as demonstrated by RT-PCR and ELISA analyses, respectively (Fig. 6b-c). This result is consistent with previous reports indicating that the depletion of UBE2N in Treg reduces IL-10 expression ^32^. TGF-β, another cytokine secreted by Tregs, remained unchanged (Fig. 6b).

**Fig. 6:**
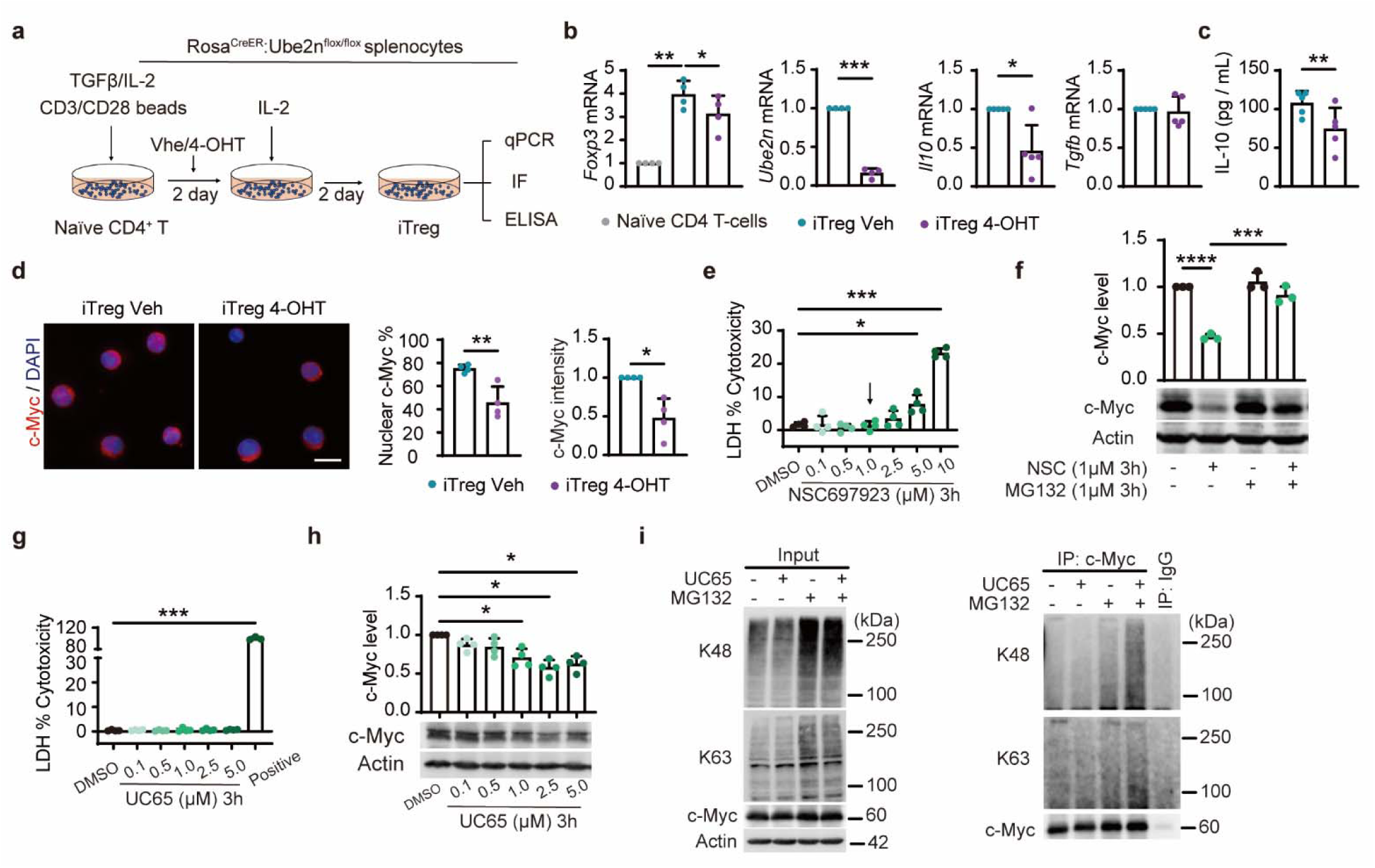
UBE2N regulates c-Myc at the protein level. **a**, Schematic of iTreg culture. **b**, Relative mRNA expression of Foxp3, Ube2n, Il10, and Tgfb in iTregs 4 days after induction. Gapdh is used for internal control. *n* = 4-5 / group, One-way ANOVA and Dunnett (Foxp3), and *t* - test. **c**, ELISA analysis of IL-10 in cell culture conditioned medium. *n* = 5 / group, *t* - test. **d**, Representative immunostaining of c-Myc (red) and DAPI nuclear staining (blue) in iTregs. Scale bar = 10 μm. Quantification of the percentage of nuclear and whole cell c-Myc intensities. *n* = 80 cells from 4 independent experiments / group, *t* - test. **e**, Quantification of lactate dehydrogenase (LDH) release in conditioned medium collected from Jurkat cells treated with varying doses of NSC697923 (NSC) for 3 hours. *n* = 4 / group, One-way ANOVA and Dunnett. **f**, Western blot and quantification of c-Myc in Jurkat cells treated with 1µM NSC with or without MG132 (1µM) for 3 hours. *n* = 3 / group, Two-way ANOVA and Tukey. **g-h**, Quantification of LDH release (**g**) and c-Myc protein levels (**h**) in Jurkat cells treated with varying doses of UC65 for 3 hours. *n* = 4 / group, One-way ANOVA and Dunnett. **i**, Immunoblotting for c-Myc, K63-Ub and K48-Ub. Protein lysates were collected from HEK293T cells which overexpressed c-Myc and treated with 2 µM UC65 with or without MG132 for 3 hours, followed by immunoprecipitation with an antibody against c-Myc. Actin was used for loading control. Data are shown as mean ± SD unless otherwise specified. *p < 0.05, **p < 0.01, ***p < 0.001, ****p < 0.0001.

As a transcription factor, activated c-Myc is mainly expressed in the nucleus. Quantitative immunofluorescence image analysis showed that both the total c-Myc and the percentage of nuclear c-Myc were reduced in UBE2N-deficient iTregs (Fig. 6d). To verify the link between UBE2N and c-Myc, we utilized human Jurkat T-cells and pharmacological agents to inhibit UBE2N. Jurkat cells were treated with varying doses of the small molecule UBE2N inhibitor NSC697923 (NSC) for 3 hours. Lactate dehydrogenase (LDH) analysis of the cell culture conditioned medium showed that 5 µM NSC significantly increased cytotoxicity, whereas 1 µM NSC was not toxic (Fig. 6e) and hence was chosen for further study. Western blotting (WB) show that c-Myc was markedly decreased following 3 hours of treatment with 1 µM NSC; proteasome inhibitor MG132 blocked c-Myc downregulation (Fig. 6f). To further validate this finding, we treated Jurkat cells with another UBE2N inhibitor, UC-764865 (UC65), which, compared to NSC, has low cytotoxicity and high specificity ^43^. As expected, no cytotoxicity was observed following 3 hours treatment of up to 5 µM UC65 (Fig. 6g). Consistently, UC65 significantly reduced c-Myc protein levels in Jurkat cells (Fig. 6h). These results indicate that UBE2N is required for the maintenance of c-Myc expression and protein stability.

We predicted that UBE2N regulates c-Myc stability via regulation of c-Myc ubiquitination. To test this idea, HEK293T cells were used to overexpress c-Myc and performed ubiquitination analyses by immunoprecipitation. We verified that UC65 decreased c-Myc and co-treatment of UC65 and the proteasome inhibitor MG132 prevented c-Myc downregulation accompanied by accumulation of c-Myc-K48-Ub (Fig. 6i). These data indicate that UBE2N maintains c-Myc protein stability via negative regulation of K48Ub-mediated proteasome degradation and positive regulation of K63-Ub either directly on c-Myc or indirectly on other regulators.

## Discussion

Our work demonstrates that Treg-targeted deletion of Ube2n in adult animals induces potent and long-lasting antitumor immunity. We also show that such a benefit is preserved after adoptive cell transfer and requires the presence of CD4^+^ T-cells along with CD8^+^ cytotoxic T-cells. We identified c-Myc as a critical downstream effector of UBE2N. In contrast to what is observed with developing animals ^32^, temporally induced deletion of UBE2N in Treg of developed animals has no apparent risk of autoimmunity. The developmentally distinct phenotypes caused by UBE2N loss at different stages of life are in line with data demonstrating that c-Myc is essential for Treg development but dispensable for the maintenance of existing mature Tregs ^28^. Presumably, Tregs developed during early life safeguard from adverse autoimmune reactions later in life, and the functions of these cells are not affected by Ube2n/c-Myc loss. Future studies are needed to confirm this possibility.

We demonstrate that both genetic and pharmacological inhibitions of UBE2N decrease c-Myc. By using in vitro iTreg, Jurkat cells, or HEK293T cells, we demonstrate that the inhibition of UBE2N leads to decreased c-Myc K63-Ub and increased degradation through a K48-Ub-mediated process. In agreement with our data, K63-Ub is required for c-Myc oncogenic activity ^44^. Nearly a dozen E3 ligases (e.g. HECT9 and TRAF6) have been linked to c-Myc ubiquitination ^44,45^. Our study identifies UBE2N as an essential E2 ligase regulating c-Myc function in Tregs. In addition, UBE2N can regulate c-Myc through indirect mechanisms. In prostate cancer cells, UBE2N promotes c-Myc expression through the Axin1/β-catenin pathway ^46^. UBE2N may also act through TRAF6 to regulate c-Myc. In live cancer, TRAF6 increases c-Myc stability through HDAC3 ^45^. On the other hand, TRAF6 ubiquitinates c-Myc and represses its transcriptional activity in myeloid leukemia ^47^. Mechanistically, how UBE2N modulates c-Myc expression and activation in Tregs in the context of cancer requires further study.

Like UBE2N, c-Myc is overexpressed in many cancers, where it regulates cell proliferation, differentiation, metabolism, and survival ^48,49^. Loss of c-Myc in Tregs during early life results in early-onset of autoimmune disorders associated with mitochondrial oxidative metabolism ^28^. Elevated c-Myc compromises Treg survival and function by inducing excessive glycolysis ^50,51^. Thus, both UBE2N and c-Myc are favorable therapeutic targets to deter cancer cell-intrinsic processes and Treg-mediated immune suppression. Pharmacological inhibitors are being developed for UBE2N and c-Myc. In fact, several small molecule inhibitors of c-Myc have shown benefits in preclinical models of various cancers with the prospect of synergy with immune checkpoint inhibitors ^49^. Specifically, Omomyc, a dominant negative Myc peptide, has achieved a clinical success in treating several solid cancers ^52^. UBE2N inhibitors such as UC-764865 have shown preclinical efficacy in treating myeloid leukemia ^43^. With the possibility of Treg-targeted drug conjugates ^53^, the findings of this study may aid the development of new strategies for therapeutic targeting.

While our data pinpoint c-Myc as a prominent mediator of UBE2N function, it is highly likely that other transcription factors are impacted by UBE2N loss. In this regard, the transcription program of FOXM1, which is both a transcriptional target and a regulator of c-Myc, is the top most downregulated in pretreatment TIL Tregs of melanoma patients who respond to ICB. Thus, the observed Treg dysfunction is likely a result of the cumulative effects of multiple gene regulators in both murine and human melanoma. In vivo, Foxp3 mRNA and protein expression levels are not significantly affected by Ube2n loss, which is in line with previous findings ^32^. However, our in vitro studies show that Ube2n deletion resulted in about 30% decrease of Foxp3 mRNA in iTreg. Additionally, like c-Myc, Foxp3 is subject to posttranslational modifications, including phosphorylation and ubiquitination ^54^. Foxp3 transcriptional activity could be altered at the posttranslational level. Treg function may also be indirectly affected by the elevated expression of the inflammatory cytokines such as IFN-γ which not only promotes cytotoxic T-cell activity but may also induce Treg fragility ^55^. Paradoxically, ICB therapies can inadvertently increase Treg activity, leading to a therapeutic resistance ^15–17^.

Lastly, our data demonstrate that the anticancer benefit of UBE2N loss in Tregs is also observed in a colon cancer model, suggesting a broad relevance of this pathway. In summary, the findings of this work underscore the feasibility of Treg-targeted strategies to boost antitumor immunity with a minimal risk of adverse autoimmune reactions. Our data highlight the UBE2N/c-Myc signaling axis as a potential therapeutic target for both inhibition of cancer cell-intrinsic processes and enhancement of antitumor immunity.

## Methods and materials

### Ethics statement

All animal experiments were performed in accordance to protocols (A075-22-04 and A200-18-08) approved by the Duke University Institutional Animal Care and Use Committee and followed guidelines set forth by the NIH and DOD for the Care and Use of Laboratory Animals. All animals were housed in a temperature and humidity-controlled facility with a 12 hours light/dark cycle, and food and water were available ad libitum. All treatments and analyses were performed by blinded investigators wherever feasible. All efforts were made to minimize animal suffering and the number of animals used.

### Experimental animals

C57BL/6J WT, Rosa26^Cre-ERT2^ (strain#008463), Foxp3^eGFP-Cre-ERT2^ (strain#:016961, referred to Foxp3^Cre-ERT2^), and Rag1-knockout (strain#034159) mice were obtained from the Jackson Laboratory (Bar Harbor, ME). Ube2n^fl/fl^ mice were kindly gifted by Dr. Shao-Cong Sun (University of Texas MD Anderson, Houston, TX, USA) under the permission of Dr. Shizuo Akira (Osaka University, Osaka, Japan)^56^. Rosa26^Cre-ERT2^.Ube2n^fl/fl^ and Foxp3^Cre-ERT2^.Ube2n^fl/fl^ mice were generated by crossbreeding. The depletion of Ube2n in Tregs was induced by intraperitoneal injection of tamoxifen 100 mg/kg daily for 3-4 consecutive days or 1 mg/mice every 2-3 days for pre and post-tumor inductions, respectively. 8 to12 week-old animals of both sexes were assigned randomly to different treatment groups. All assessments were performed by investigators who were blinded to experimental group assignments.

### Mouse tumor model

Two to three-month-old mice were used for tumor implantation. B16-F10 cells (1x10^5^) or MC38 cells (2x10^6^) suspended in 100 µL PBS: matrigel (2:1) were injected subcutaneously into the right and left flanks of each animal. Tumor size and body weight were measured every 3 days for 2-4 weeks or longer. Tumor images were analyzed using NIH ImageJ to calculate tumor sizes. Mice were euthanized and tissue samples were collected for endpoint analysis.

### Cell culture

The B16-F10 murine melanoma, MC38 murine colon carcinoma, Jurkat human T-lymphocyte, and HEK293T cell lines were purchased from ATCC via the Duke Cell Culture Facility. B16-F10, MC38, and HEK293T cells were cultured in DMEM (ThermoFisher, Cat# 11965092) supplemented with 10% FBS (R&D, Cat# S11150H) and 1X Antibiotic-Antimycotic (ThermoFisher, Cat# 15240096). Jurkat cells were cultured in RPMI 1640 (ThermoFisher, Cat# 11875093) supplemented with 10% FBS and 1X Antibiotic-Antimycotic. Cells were maintained in a 37°C incubator with 5% CO_2_, and pathogen-negative cells were used for experiments.

### iTreg induction

Naïve CD4 T-cells were isolated from the spleens of 4-week-old Rosa26^Cre-ERT2^.Ube2n^fl/fl^ mice using the naïve CD4 T-Cell Isolation Kit (Miltenyi, Cat#130-104-453), following the manufacturer’s instructions. Cells were resuspended at 5x10^5^ cells/mL in RPMI 1640 complete medium with 10% FBS, 1X Antibiotic-Antimycotic, 10 mM HEPES, and 0.1 mM non-essential amino acids. Murine recombinant IL-2 (100 U/mL; PeproTech, Cat# 212-12), murine recombinant TGF-β (5 ng/mL; BioLegend, Cat# 763102), and 25 µL Dynabeads™ Mouse T-Activator CD3/CD28 (ThermoFisher, Cat# 11456D) were added for T-cell expansion and induction. After 48 hours of incubation, an equal volume of complete medium with 2000 U/mL of recombinant IL-2 was added for an additional 2 days of culture ^41,42^. Cells were treated with (Z)-4-Hydroxytamoxifen (1 µM, Selleckchem, Cat#S8956) starting on day 1 to deplete UBE2N in iTregs. Subsequently, the cells and the conditioned medium were collected for iTreg validation and endpoint analysis.

### Adoptive T-cell transfer

Single-cell suspensions were prepared from the spleens of tumor-cleared Ube2n^Treg-KO^ mice or age-matched WT naïve mice. Pan-T-cells were isolated using the T-Cell Isolation Kit II (Miltenyi, Cat# 130-095-130) following the manufacturer’s instructions. Cells were resuspended in PBS (1x10^6^ cells/100 µL/injection) and intravenously injected into Rag1-KO mice within 10 min after B16 tumor implantation. For CD8^+^ T-cell enrichment, Pan-T-cells were further incubated with biotin-conjugated anti-CD4 antibody, followed by negative selection with anti-biotin beads. CD8^+^ T-cells were resuspended in PBS at 5x10^5^ cells/100 µL for intravenous injection.

### Flow cytometry

Blood samples were collected from the heart of deeply anesthetized mice which were then perfused with 30 mL of 0.9% NaCl and sacrificed. Tumors were dissociated using 1 mg/mL collagenase type 4 (Worthington, Cat# LS004188) in RPMI 1640 supplemented with 50 µg/mL DNase I (MP Biological, Cat# 190062) at 37°C for 45 min, with pipetting every 5 min. Samples were then filtered through a 40 µm strainer and centrifuged. The cell pellet was resuspended in 1 mL of RBC lysis buffer (BioLegend, Cat# 420301) and incubated for 5 min at room temperature to remove red blood cells. Single-cell samples from blood, spleen, thymus, or tumor were resuspended in FACS buffer (1x PBS supplemented with 1% FBS and 0.5 mM EDTA). Ghost Dye™ Violet 510 (1 µL/test, TONBO, Cat# 13-0870-T500) was used for live and dead cell labeling, followed by fluorophore-conjugated primary antibodies for cell surface protein labeling. For cytoplasmic protein staining, the eBioscience™ Intracellular Fixation & Permeabilization Buffer Set (ThermoFisher, Cat# 88-8824-00) was used. For Foxp3 staining, the eBioscience™ Foxp3 Transcription Factor Staining Buffer Set (ThermoFisher, Cat# 00-5523-00) was used according to the manufacturer’s instructions. Appropriate isotype controls were used per the manufacturers’ instructions (BioLegend). Fluorochrome compensation was performed with single-stained spleen cells using fluorophore-labeled CD4 or CD8 antibodies. Flow cytometry was performed on a BD FACSCanto-II (BD Biosciences, San Jose, CA) according to the manufacturer’s instructions. Data were analyzed using FlowJo software (FlowJo 10.0, Ashland, OR). Conjugated primary antibodies were purchased from Biolegend: CD45-APC (Clone# 30-F11, Cat# 103112); CD11b-PE/Cyanine7 (Clone# M1/70, Cat# 101216); CD11c-APC/Cyanine7 (Clone# N418, Cat# 117324); F4/80-PE (Clone# BM8, Cat# 123110); Ly6G-Pacific Blue (Clone# 1A8, Cat# 127611); Ly6C-PerCP/Cyanine5.5 (Clone# HK1.4, Cat# 128011); CD3-APC/Cyanine7 (Clone# 17A2, Cat# 100222); CD4-Pacific Blue (Clone# RM4-5, Cat# 100531); CD8-PE/Cyanine7 (Clone# 53-6.7, Cat# 100722); CD25-PerCP/Cyanine5.5 (Clone# PC61, Cat# 102029); Foxp3-Alexa Fluor® 488 (Clone# MF-14, Cat# 126405); IFN-γ -PE (Clone# XMG1.2, Cat# 505808).

### Immunofluorescence and image analysis

Tumors were frozen in OCT for cryosectioning, and 10 µm tumor sections were used for immunofluorescence staining as previously described ^57^. In brief, sections were fixed in methanol for 15 min at -20°C. After washing with PBS, sections were blocked with 10% donkey serum in 0.05% Tween-20 PBS for 1 hour at room temperature. Primary antibodies were diluted in 1% donkey serum in 0.05% Tween-20 PBS at optimized concentrations and incubated with shaking overnight at 4°C, followed by incubation with conjugated secondary antibodies. The primary antibodies used in this study include the following: Alexa Fluor® 647 anti-mouse CD3 antibody (Biolegend, Clone# 17A2, Cat# 100209); Alexa Fluor® 488 anti-mouse CD4 antibody (Biolegend, Clone# RM4-5, Cat# 100532); Purified Rat Anti-Mouse CD8b.2 antibody (Biolegend, Clone# 53-5.8, Cat# 553038); CD45-APC (Biolegend, Clone# 30-F11, Cat# 103112); Anti-MLANA antibody (Sigma Aldrich, Cat# HPA048662); c-Myc (Proteintech, Cat#10828-1-AP; Rosemont, IL). Secondary antibodies were purchased from Themo Fisher Scientific. All the secondary antibodies were diluted 1:1000. The images were acquired on an Olympus IX73 microscope (Olympus, Tokyo, Japan) with a UPLSAPO 10 or 20× objective. Image analyses were performed on 4-8 randomly selected microscopic fields within 1 mm from tumor periphery. Three sections were assessed for each tumor. Images were coded and analyzed using the Image-Pro Plus software (Media Cybernetics, Rockville, MD) and/or ImageJ/Fiji (National Institutes of Health, Bethesda, MD) by an investigator blinded to experimental groups.

### Real-time PCR

Total RNA was extracted from tumor samples and cells using the TRIzol reagent (Invitrogen, Cat# 15596018; Carlsbad, CA). RNA (1 µg) was used to synthesize cDNA using the iScript™ Reverse Transcription Supermix System followed by RT-PCR according to the manufacturer’s protocols (Bio-Rad, Cat# 1708840; Hercules, CA). The reverse transcription program was 25°C for 5 min, 46°C for 20 min, 95°C for 1 min, and hold at 4°C. PCR was performed on the Applied Biosystems™ StepOne™ Real-Time PCR System (Thermo Fisher Scientific, Cat# 4376357; Pittsburgh, PA) using corresponding primers (Table S1) and 2X Universal SYBR Green Fast qPCR Mix (ABclonal, Cat# RK21203; Woburn, MA). The real-time PCR program was 95°C for 15 min, followed by 40 cycles of 94°C for 20 s, 59°C for 30 s, and 72°C for 30 s. The melting curve was from 50°C to 92°C, with readings every 0.2°C, hold for 2 s, and incubation at 8°C. Cycle threshold (Ct) values were normalized to Gapdh for internal control.

### ELISA

iTregs were plated in a 12-well plate at 5x10^5^ cells per well. Conditioned medium from iTreg cultures was collected on day 4 after cytokine induction. Mouse IL-10 levels were measured using a commercial ELISA quantification kit according to the manufacturer’s instructions (BioLegend, Cat# 431411).

### Lactate dehydrogenase assay

Jurkat cells were plated in a 24-well plate at 1 x 10^6^ cells per well and treated with varying concentrations of NSC697923 (MCE, Cat# HY13811) and UC-764865 synthesized as described ^43^. Cell culture conditioned medium was collected 3 hours after treatment and used for lactate dehydrogenase assay (Invitrogen, Cat# C20300) according to the manufacturer’s protocol.

### Western blot

For protein collection, cells were lysed on ice for 15 min using RIPA buffer with Halt™ Protease Inhibitor (Thermo Fisher Scientific, Cat# 78429), Halt™ Phosphatase Inhibitor (Thermo Fisher Scientific, Cat# 78420), 10 mM N-ethylmaleimide (Selleckchem, Cat# S3692; Houston, TX), and 10 mM iodoacetamide (IAA, UBPBio, Cat# P1052). Western blots were performed using standard SDS-polyacrylamide gel electrophoresis (PAGE) with 30 µg of total protein loaded per lane. Immunoreactivity was semi-quantitatively measured by gel densitometric scanning and analyzed using ImageJ software. The primary antibodies used in this study include c-Myc (1:1000, Proteintech, Cat#10828-1-AP; Rosemont, IL) and β-actin (1:3000, Cat# 4970; Cell Signaling Technology, Danvers, MA). K63-linkage specific polyubiquitin (Cells Signaling, Cat#5621); K48-ubiquitin (Millipore, Cat#051307).

### Single-cell RNA library preparation and sequencing

Tumors were excised 14 days after engraftment, manually dissociated, and digested with 1 mg/mL collagenase type 4 (Worthington, Cat# LS004188) in RPMI 1640 with 50 µg/mL DNase I at 37°C for 45 min, with pipetting every 5 min. Tumor-infiltrating lymphocytes (TILs) were isolated using CD4/CD8 MicroBeads (Miltenyi, Cat# 130-116-480) according to the manufacturer’s protocols. Single cell suspensions from 3 animals per group were pooled, centrifuged, and washed twice in PBS containing 0.04% RNase-Free BSA (weight/volume). An estimate of 1.2x10^6^ cells/group were sent to Duke Molecular Genomics Core for single cell library construction. Cells were size selected and quantified (Cellometer, Nexcelom - Lawrence, MA) and normalizing to 1x10^3^ cells/ul. Libraries were tittered to ∼10,000 cells per library and resuspended in master mix combined with gel beads carrying the Illumina TruSeq Read 1 sequencing primer, barcode, unique molecular identifier and a poly-dT primer or gene specific probes for RT (10xGenomics, Pleasanton, CA). Full length cDNAs were cleaned and assayed (Agilent TapeStation, Santa Clara, CA) to ensure library lengths between 200-5000bp. Enzymatic fragmentation occurred prior to Illumina (San Diego, CA) sequence adapter and sample index ligation; TruSeq read 2 primers are added via End Repair, A-tailing, Adaptor Ligation, and PCR. qPCR will be used to assess P5 and P7 adapter ligation, prior to library QC assessment to ensure the libraries are sized between 400-500bp. Sequence was generated with 150bp paired end sequencing on an Illumina NovaSeq 6000 Illumina sequencing platform at 50K reads per cell.

### Single-cell RNA-sequence analysis workflow

The primary analytical pipeline for the scRNA-seq analysis followed the recommended protocols from 10X Genomics. Briefly, we demultiplexed the generated raw base call (BCL) files into FASTQ files. The CellRanger (version 7.1.0) count function was used to align the reads to the mm10 mouse reference genome and generate the cell-gene count matrix following filtering unviable cells based on low number of unique RNA molecules per cell. The filtered count matrix was read using Seurat (version 4.1.1) ^58^ on R (version 4.0.5). Seurat was used to create a Seurat object from the read count matrix. Subsequent quality control, exploratory, and differential expression analyses were also done using Seurat. Cells were further filtered based on the number of unique RNA molecules observed (minimum of 500 and maximum of 100,000), percent of counts ascribed to mitochondrial genes (maximum of 20%), the number of detected genes per cell (minimum of 250 genes and maximum of 9,000). The thresholds were used to prune out outliers based on an initial plot of these quality control metrics before filtering to exclude low quality cells and possible doublets. The counts were then normalized for sequencing depth using the SCTransform method ^59^ during which the percentage of mitochondrial counts was used as a regression variable. Cells from the various batches of sequencing were then integrated using the Seurat anchor method ^60^ using the top 3000 variable features and with SCTransform as the normalization method. Subsequently, clustering (resolution of 0.8) and UMAP were done using 35 principal components. Identification of cell types as well as the choice of optimal clustering parameters were guided by the localization of cell type-specific genes. Marker genes used for each cell type have been shown in the corresponding figures in this manuscript. Sub-clustering of immune cells (resolution of 0.7) and UMAP were done using 20 principal components. For differential expressed gene (DEG) analysis, a false discovery rate (also known as adjusted p-value) of less than 0.05 and a log2 fold-change of 0.5 were used as thresholds to call significantly dysregulated genes. EnrichR ^37–39^ (version 3.2) was used for gene ontology enrichment analysis. GO terms and associated gene lists used for calculating module score were obtained from Gene Ontology Browser tool from the Mouse Genome Informatics (https://www.informatics.jax.org/vocab/gene_ontology/). scRNA data are deposited to NCBI (accession # GSE256255).

For human data, the originally described G7 cluster of the publicly available single-cell RNAseq data (GSE120575) ^40^ was subject to UMAP analysis at clustering with resolution of 0.2 using 10 principal components. The pre-treatment baseline Tregs populations of subcluster 1 were used for DEG analysis comparing responder to non-responder and subsequent EnrichR analysis, as described above. Heatmap was generated with the putative c-Myc target gene list obtained via EnrichR of DEGs with adjusted p-values <0.05.

### Statistical analysis

Sample sizes for animal studies were determined based on pilot studies or the literature. Results are presented as mean ± standard deviation (SD). GraphPad Prism software (version 7.0.0, La Jolla, CA) was used for statistical analyses. The Student’s *t* - test was used for comparisons of two groups with continuous variables that follow normal distributions. The Mann-Whitney U rank-sum test was used for continuous variables with non-normal distributions. Differences in means among multiple groups were analyzed using the Kruskal-Wallis test followed by Dunn’s test, or one-way or two-way analysis of variance (ANOVA) followed by Dunnett’s test (all conditions compared with a specific group) or Bonferroni or Tukey’s multiple-comparison tests (comparisons between all conditions). Differences in means across groups with repeated measurements over time were analyzed using repeated measures ANOVA, followed by post hoc Tukey or Bonferroni tests. Spearman rank correlation analyses were used to test correlations between data sets with non-normal distributions. In all analyses, p < 0.05 was considered statistically significant.

## Supporting information

Table S1 RT-PCR Primers

## Acknowledgements

This study was supported by funding from DOD (W81XWH-22-1-1061 ME210022), NIH/NIAMS (R01AR068991) to JZ. We thank Dr. Shizuo Akira of Osaka University and Dr. Shao-Cong Sun of University of Texas MD Anderson for providing the Rosa^CreER^.Ube2n^fl/fl^ mice, as well as Drs. Xiaoping Zhong, Edward Miao, and Fang Li (Duke University, Durham, NC) for sharing their resources and expertise of Foxp3^Cre-ERT2^ mice, Rag1-KO mice, and M38 cells, respectively. We would also like to thank the staff of the Duke Molecular Genomics Core for generation of the single cell sequencing libraries.

