## Supplemental Text and Figures S1-S4 for "Inhibition of UBE2N in regulatory T-cells boosts immunity against cancer"

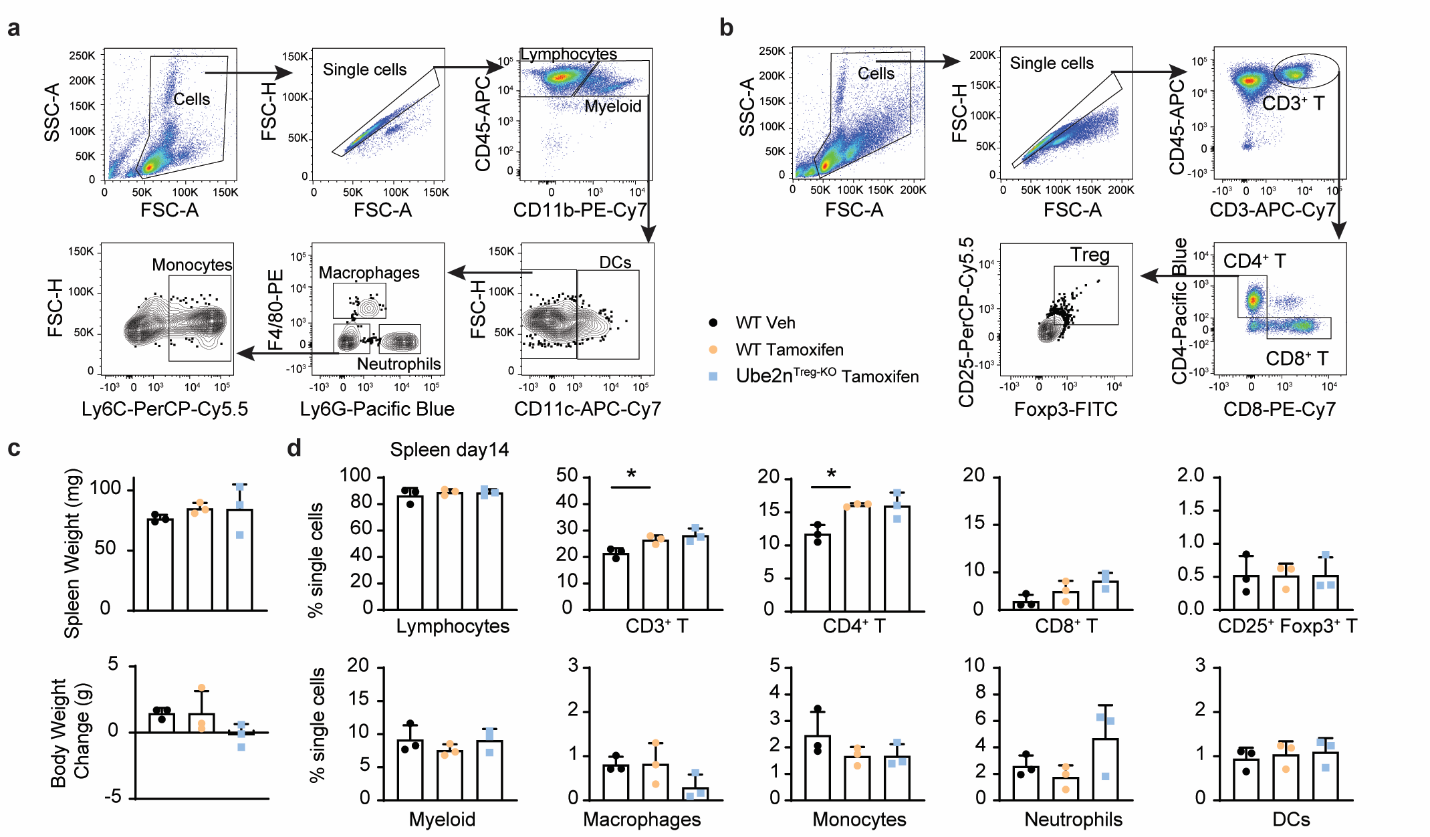

**Fig. S1:** **Treg-specific deletion of UBE2N did not affect the splenocyte population.** **a-b**, Gating strategy for spleen Lymphocytes (CD45^+^CD11b^-^), CD3^+^ T-cells (CD45^+^CD11b^-^CD3^+^), CD4^+^ T-cells (CD3^+^CD4^+^CD8^-^), CD8^+^ T-cells (CD3^+^CD4^-^CD8^+^), Tregs (CD4^+^CD25^+^Foxp3^+^), myeloid cells (CD45^+^CD11B^+^), macrophages (CD11b^+^CD11c^-^Ly6G^-^F4/80^+^), neutrophils (CD11b^+^CD11c^-^Ly6G^+^F4/80^-^), monocytes (CD11b^+^CD11c^-^Ly6G^-^F4/80^-^Ly6C^+^), and dendritic cells (DCs) (CD11b^+^CD11c^+^). **c**, Spleen weight (upper) and body weight change (bottom) 14 days after tamoxifen treatment. **d**, Quantification of spleen lymphocytes, CD3^+^ T-cells, CD4^+^ T-cells, CD8^+^ T-cells, Tregs, myeloid cells, neutrophils, monocytes, and DCs 14 days after tamoxifen treatment. *n* = 3 mice / group, One-way ANOVA and Dunnett’s. Data are shown as mean ± SD unless otherwise specified. *p < 0.05.

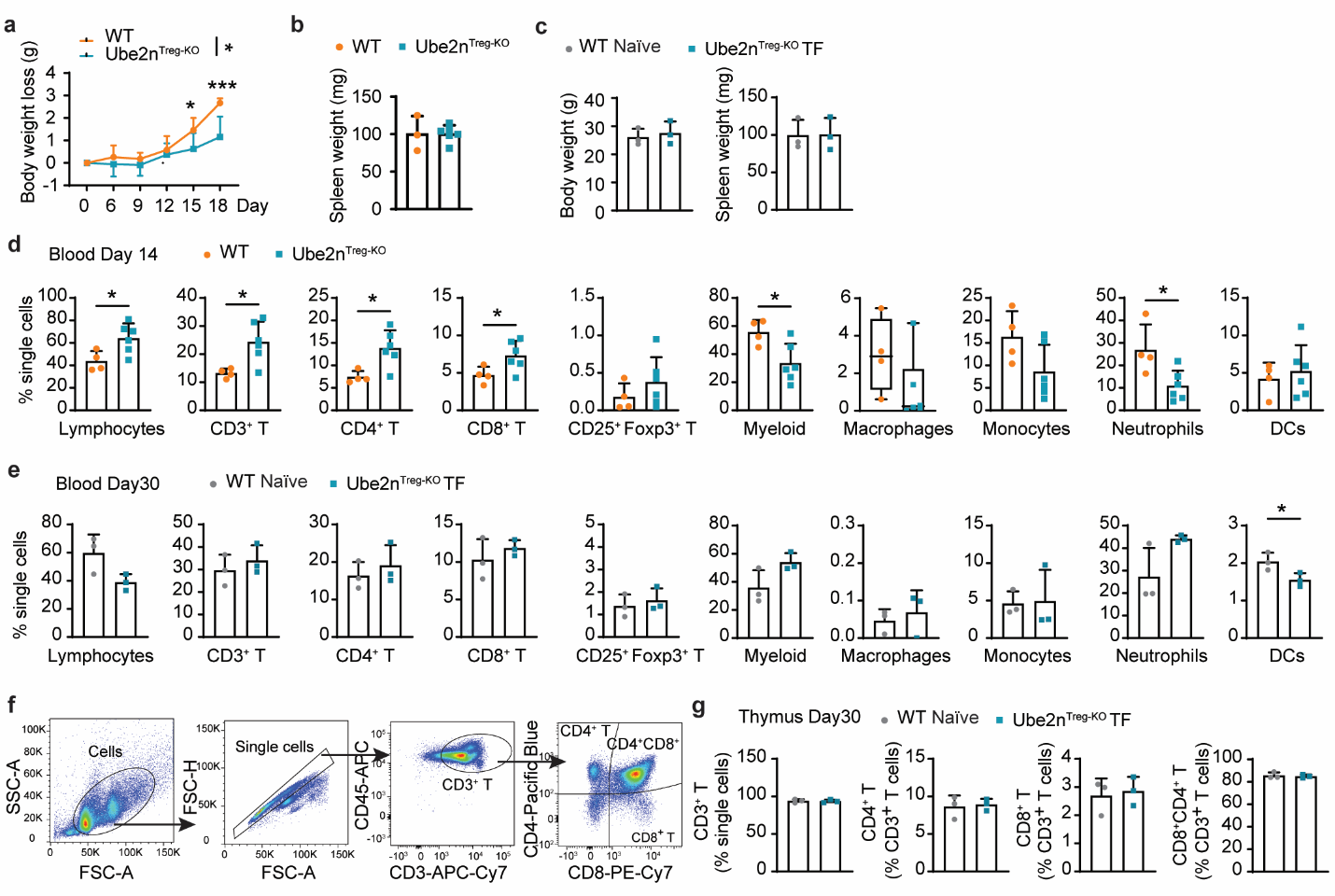

**Fig. S2:** **Immune cells returned to normal levels in tumor-eradicated Ube2n^Treg-KO^ TF mice at 1 month after tumor implantation.** **a**, Body weight loss of tumor-bearing mice 18 days after tumor implantation. WT *n* = 5 mice, Ube2n^Treg-KO^ *n* = 9 mice, Two-way ANOVA and Bonferroni. **b**, Spleen weight 14 days after tumor implantation. *n* = 3-6 mice/group, *t* - Test. **c**, Body weight (left) and spleen weight (right) of WT B16^-^ (age-matched nontumor-bearing mice) and Ube2n^Treg-KO^ TF (KO tumor-free mice) 30 days after tumor implantation. *n* = 3 mice/group, *t*-Test. **d-e**, Quantification of blood immune cells14 days (**d**) and 30 days (**e**) after tumor implantation. Day 14, *n* = 4-6 mice / group, *t* - Test. Day 30, *n* = 3 mice/group, *t* - Test. **f**, Gating strategy for thymus CD3^+^ T-cells (CD45^+^ CD3^+^), CD4^+^ T-cells (CD3^+^CD4^+^CD8^-^), CD8^+^ T-cells (CD3^+^CD4^-^CD8^+^), and CD4^+^CD8^+^ T-cells (CD3^+^CD4^+^CD8^+^). **g**, Quantification of thymus lymphocytes 30 days after tumor implantation. *n* = 3 mice / group, *t* - Test. Data are shown as mean ± SD unless otherwise specified (**d**: Macrophages; Data are presented as a box-and-whisker plot, with bounds from 25^th^ to 75^th^ percentile, median line, and whiskers ranging from minimum to maximum value). *p < 0.05, ***p < 0.001.

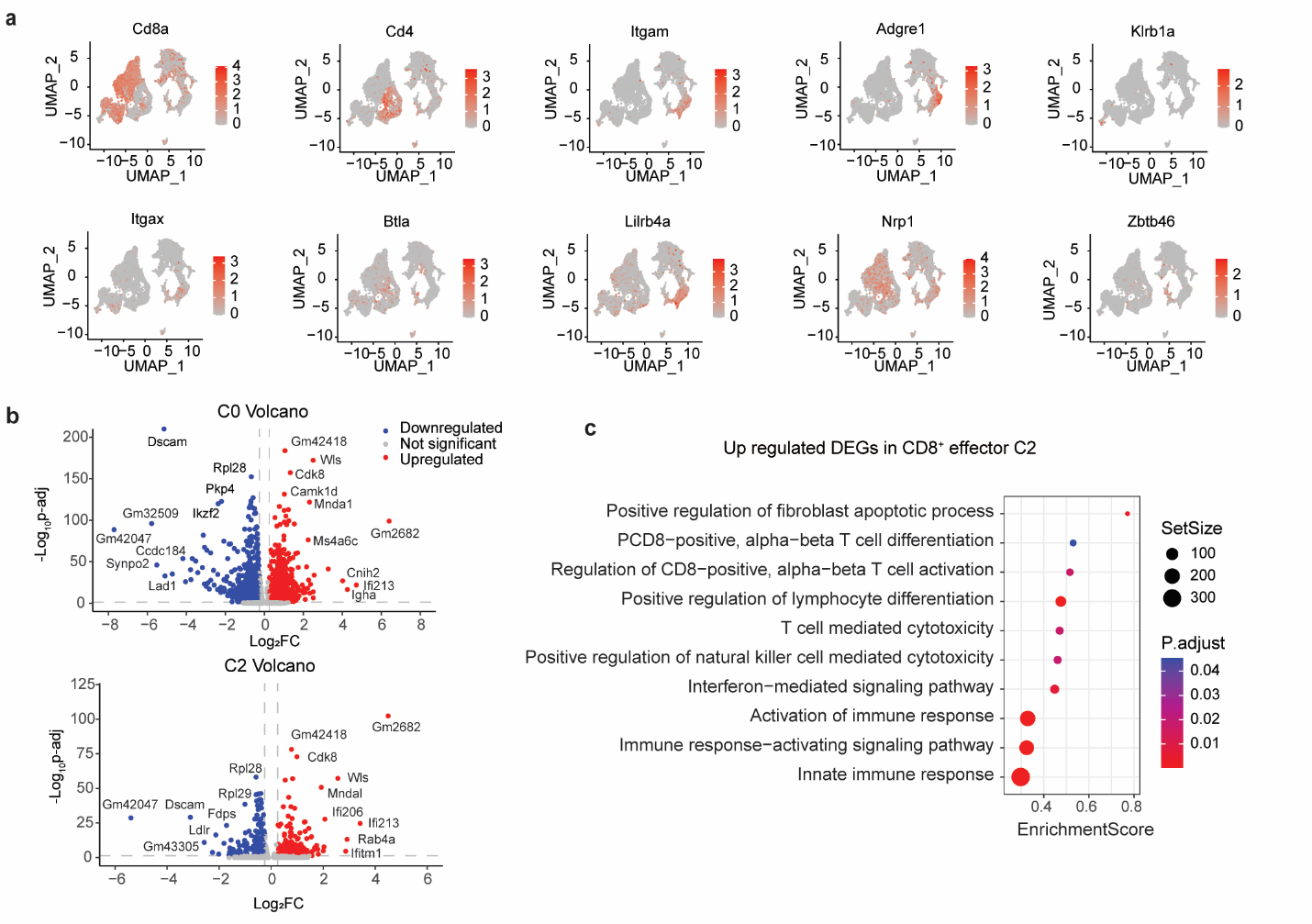

**Fig. S3: Immune cell identification and GO of Cluster 2. a**, Marker gene expression profile plot by integrated UMAP. **b**, Volcano plot DEGs in WT vs. Ube2n^Treg-KO^ in CD8^+^ effector/exhaustion cells (cluster 0, upper) and CD8^+^ effector cells (cluster 2, bottom). **c**, GO enrichment result of upregulated DEGs of WT vs. Ube2n^Treg-KO^ in CD8^+^ effector cells (cluster 2).

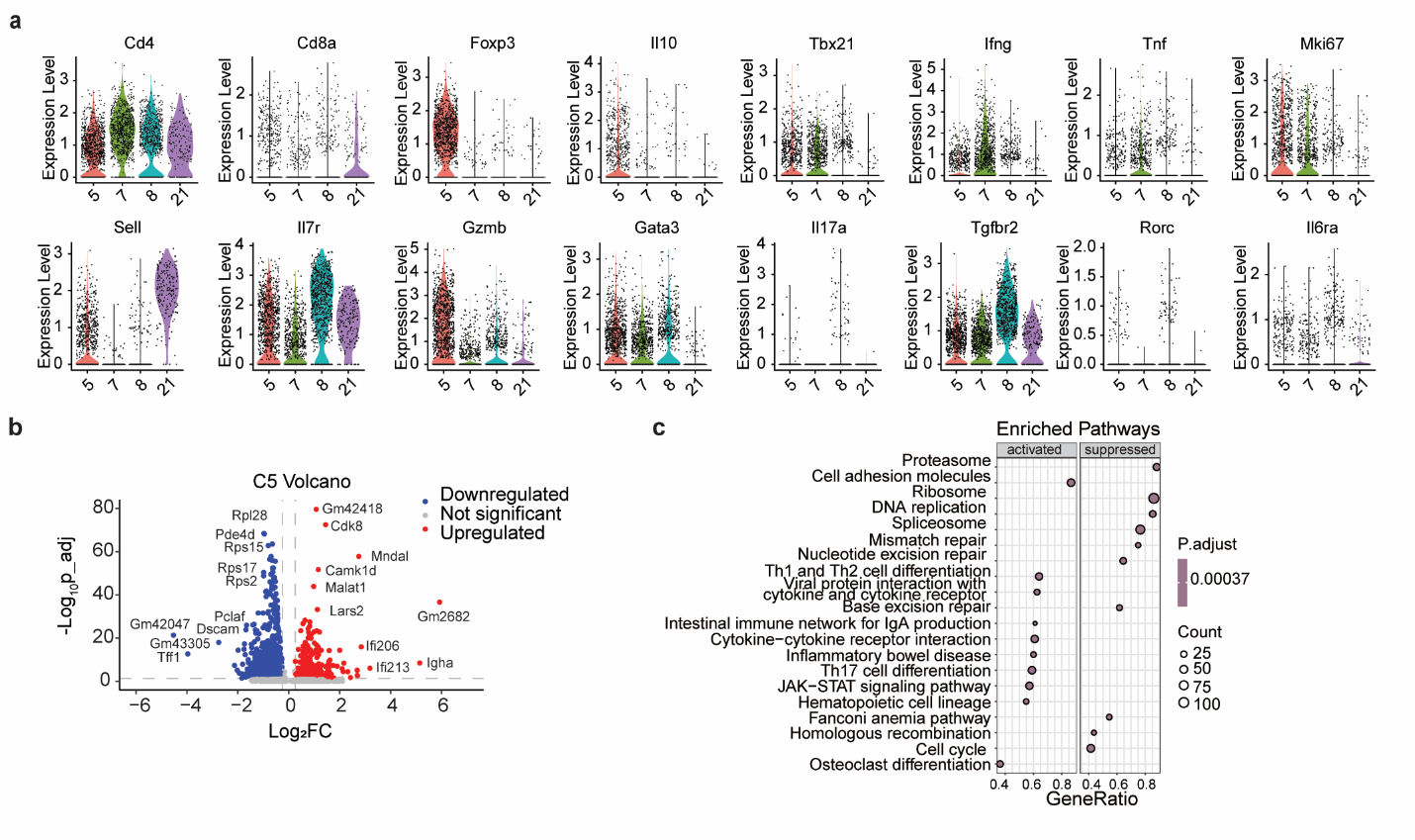

**Fig. S4:** **CD4^+^ T cell subtype identification and KEGG of Tregs.** **a**, Violin plots of marker gene expression in CD4^+^ Tregs. **b**, Volcano plot DEGs in Tregs WT vs. Ube2n^Treg-KO^ Tregs (cluster 5). **c**, KEGG enrichment result of DEGs of WT vs. Ube2n^Treg-KO^ Tregs (cluster 5).
